# Tandem Duplicate Genes in Maize are Abundant and Date to Two Distinct Periods of Time

**DOI:** 10.1101/238121

**Authors:** Thomas J. Y. Kono, Alex B. Brohammer, Suzanne E. McGaugh, Candice N. Hirsch

## Abstract

Tandem duplicate genes are proximally duplicated and as such occur in the same genomic neighborhood. Using the maize B73 and PH207 *de novo* genome assemblies, we identified thousands of tandem gene duplicates that account for ~10% of the genes. These tandem duplicates have a bimodal distribution of estimated ages corresponding to known periods of genomic instability. Tandem duplicates had a number of associated features that suggest origins in nonhomologous recombination based on smaller size distribution and higher rate of containing LTRs than non-tandem duplicates. Within relatively recent tandem duplicate genes, ~26% appear to be undergoing degeneration or divergence in function from the ancestral copy. Our results show that tandem duplicates are abundant in maize, arose in bursts throughout maize evolutionary history under multiple potential mechanisms, and may provide a substrate for novel phenotypic variation.

## INTRODUCTION

Gene duplications provide a mechanism through which functional novelty may arise. Many protein coding genes in eukaryotes are part of large families of genes with related function and are consistent with origins in gene duplication (Rubin 2000). The pattern of duplicate gene distribution across the genome can have important consequences for the evolutionary fate of duplicate genes. For example, genes that are duplicated in tandem (proximal in the genome), and in the same genomic neighborhoods and potentially have shared regulatory elements, and thus may diverge differently than dispersed duplicates.

The initial impact of gene duplication on phenotypes likely occurs via gene dosage effects. In many cases, the sudden change of gene product concentration has deleterious effects on the physiology of the organism and will be selected against (Innan and Kondrashov 2010). In some cases, however, the increased gene expression may be beneficial, and there will be selection to maintain the duplication (e.g., tandem duplications conferring soybean cyst nematode resistance at Rhg1 (Cook et al. 2012)). Over evolutionary time, the fate of tandem duplicate genes is less straightforward than simply either retention or purging. Mutations in the regulatory regions or mutations in the coding sequence, may cause the duplicates to be expressed in different tissues or may engender non-redundant functional roles (Flagel and Wendel 2009). Several models such as the “duplication-degeneration-complementation” model or the “escape from adaptive conflict” model have been used to describe these scenarios as possible subfunctionalization outcomes for tandem duplicates (Innan and Kondrashov 2010).

Many studies of tandem duplicates have focused on specific gene families. For instance, resistance (R) gene clusters and ribosomal gene clusters have been extensively studied tandem duplicates (Hill et al. 1977; Anderson and Roth 1981; Leister 2004). In addition, a number of classical tandem duplicates have been identified in mapping studies, owing to a large phenotypic impact. For example, the r locus in maize was determined to be tandem duplicated genes by crossing and observation of recombination frequency (Dooner and Kermicle 1971). Other examples of classical tandem duplicates that have been discovered in maize and contribute to phenotypic variation include the *White Cap* (*Wc*) locus (Tan et al. 2017), the *anthocyaninless1 (a1)* locus (Laughnan 1952), and the P locus (Athma and Peterson 1991) which all influence to grain color variation, the MATE1 locus that contributes to aluminum tolerance (Maron et al. 2013), and the *Tunicate1* (*Tu1*) locus that results in the characteristic phenotype of pod corn in which the glum covers the kernel (Han et al. 2012; Wingen et al. 2012).

Identification of tandem duplicates through phenotypic analysis can bias the understanding of genome-wide rates, evolutionary impacts, and potential phenotypic impacts of tandem duplicates within the genome. In contrast, genome-wide studies to identify tandem duplicates can identify duplicates in a way that does not condition on the duplicate visibly altering a phenotype. On a genome-wide basis, tandem duplicates may be identified from blast similarity in long sequencing reads (Dong et al. 2016), optical maps (Mak et al. 2016), or by orthologous searches of all genes within an assembled reference genome (Cannon et al. 2004). Alternatively, *de novo* assemblies of multiple individuals within a same species would provide an ideal setting for high-resolution identification and analysis of variance for tandem duplicates within a species. There are few plant species with multiple *de novo* genome assemblies, and as such, genome-wide studies of tandem duplicate gene variation across multiple individuals within a species have been limited to date in plants.

While a number of tandem duplicates have been deeply characterized for their phenotypic effect and there are descriptive studies on tandem duplicate identification in plant species, there still remain a number of important questions surrounding tandem gene duplication to understanding genome evolution. Maize is an excellent model system in which to ask these questions due to the genomic resources available within the species (multiple publicly available genome assemblies) and an understanding of the evolutionary history of this species including a recent whole genome duplication event. The major questions we aim to address in this study are 1) How many genes are tandemly duplicated in the genome, and to what extent are tandem duplicates shared or private between individuals within the species?; 2) Do tandem duplicates arise during previously identified periods of genome instability?; 3) Can we decipher potential mechanisms that generate tandem duplications?; and 4) Do tandem duplicate genes in maize show a different substitution rate relative to other maize genes and other grass genes?

## RESULTS

### Tandem duplicate gene clusters are prevalent in maize genomes

To identify tandem duplicate gene clusters (i.e. proximally duplicated groups of genes) we utilized the B73 *de novo* genome assembly generated with single-molecule technology (Jiao et al. 2017) and the PH207 *de novo* short-read genome assembly (Hirsch et al. 2016). The singlemolecule technology used for the B73 genome assembly provides a high confidence assembly for evaluating tandem duplicate clusters. In assembling the PH207 genome, great attention was given to not collapse tandem duplicate gene clusters (Hirsch et al. 2016). However, by the nature of a short read assembly, the PH207 *de novo* assembly will likely have an underrepresentation of the total tandem duplicate clusters.

Putative tandem duplicates were identified and curated based on a weighted similarity metric that allowed for some interspersed genes. In total, 1,758 tandem duplicate clusters in B73 and 1,467 tandem duplicate clusters in PH207 were identified (table 1 and supplementary table 1). The total number of annotated genes in tandem duplicate clusters was 4,448 (11.3% of the total genes) in B73, and 3,788 (9.3% of the total genes) in PH207. As expected, the number of tandem duplicate clusters and the number of genes within tandem duplicate clusters was slightly lower in PH207. The B73 abundances are likely a more accurate representation of the number of tandem duplicate genes within the maize genome.

**Table 1.**
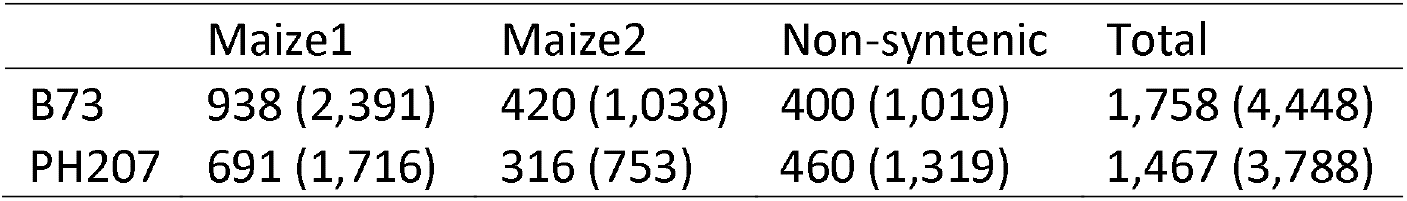
Tandem duplicate gene cluster and gene counts by subgenome in B73 and PH207. Numbers of clusters are given outside of the parenthesis and number of genes in clusters are given in parentheses.

|  | Maize1 | Maize2 | Non-syntenic | Total |
| --- | --- | --- | --- | --- |
| B73 | 938 (2,391) | 420 (1,038) | 400 (1,019) | 1,758 (4,448) |
| PH207 | 691 (1,716) | 316 (753) | 460 (1,319) | 1,467 (3,788) |

### Similar numbers of shared and private tandem duplicates are observed within species

Having access to multiple *de novo* genome assemblies within maize allowed us to determine the consistency of tandem duplicate gene cluster characteristics within the species. A similar distribution was observed between B73 and PH207 for the number of genes per cluster, and the majority of tandem duplicate genes clusters contained only two genes (fig. 1A). Within clusters, most genes were directly adjacent with no intervening genes in both B73 and PH207 (supplementary fig 1). Based on the algorithm that was used, genes were considered tandem duplicates with up to 15 intervening genes. Intervening genes were permitted to account for mechanisms that would not cause a duplicate to be directly adjacent but in the same genomic neighborhood, to allow for instances in which a gene is inserted after the duplicate event, and for possible misassembly and misannotation of genes. Only 17% of the duplicate gene clusters that were identified had an interval of greater than two intervening genes between members of the cluster. To determine if specific tandem duplicates were shared between the assemblies, we used previous information that linked the B73 and PH207 gene models (Brohammer et al. 2018). Interestingly, only about half of the B73 tandem duplicate gene clusters and 60% of the PH207 tandem duplicate clusters were shared between the two genomes (fig. 1B). This suggests that the formation and loss of tandem duplicates is an ongoing process, with new duplications arising or being lost after the divergence of B73 and PH207.

**Fig. 1.**
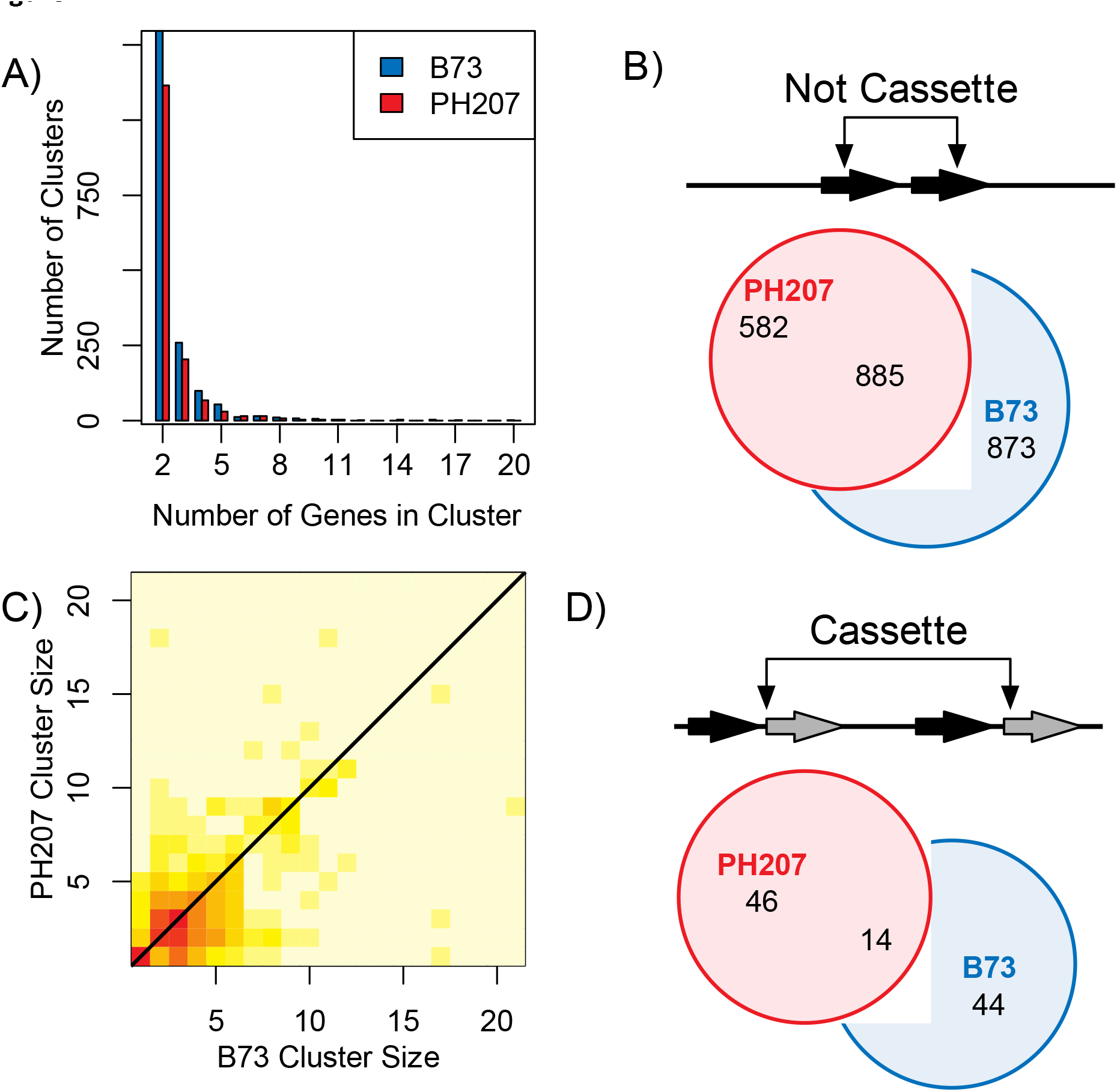
Maize tandem duplicate cluster summary. A) The distribution of cluster sizes in B73 v4 and PH207 v1 genome assemblies. B) Euler diagram of shared and private non-cassette tandem duplicate gene clusters for B73 and PH207. C) Heatmap of log of the number of instances of tandem duplicate gene cluster size relationships between B73 and PH207 (N=4,393 clusters). Color scale ranges from cream equals zero to red equals 1,257 genes in a cluster. D) Euler diagram of shared and private cassette duplicate gene clusters for B73 and PH207.

Another way that individuals can differ with regards to tandem duplicates is the number of duplicate copies within a shared tandem duplicate cluster. To determine the rate of difference in tandem duplicate copy number, we evaluated the 885 tandem duplicate clusters that were shared between the B73 and PH207 genome assembly. For the most part, tandem duplicate genes shared between B73 and PH207 exhibited similar cluster sizes as shown by strong heat along the diagonal in fig. 1C. That is, when homologous genes were part of tandem clusters in B73 and PH207, these clusters often contained similar numbers of genes. However, there was variation in tandem duplicate cluster size with a difference of up to 16 more copies in one of the genotypes compared to the other (fig. 1C).

### Cassette tandem duplication events are rare and often private

Tandem gene duplication events can occur as a single gene duplication event or in sets of genes that duplicated as a tandem cluster (i.e. Gene A-1 Gene B-1 followed by Gene A-2 and Gene B-2). Cassette tandem duplicate gene clusters likely arise from a single event in which a set of genes is duplicated simultaneously, thought, it is possible for tandem duplicate cassettes to be generated from multiple duplication events. Candidate tandem duplicate cassettes were identified from interlaced tandem duplicate gene clusters (supplementary fig. 2). Cassette duplications were rare in both of the inbred lines with only 58 and 60 cassette duplications in B73 and PH207, respectively (fig. 1D). A higher frequency of private cassette tandem duplicates was observed relative to non-cassette tandem duplicates (fig. 1B and 1D). Only 14 cassettes were shared across genotypes, which equates to approximately one-quarter of the tandem duplicate cluster cassettes in either genome. There was variation for the composition of the cassette between the two genotypes for 10 of the 14 shared cassettes. These differences may be the result of multiple duplication events in one genotype that did not occur in the other, but they are more likely the product of differential loss of a common complete duplication event.

### Tandem duplicate clusters are dispersed throughout the genome and correlate with gene density

Tandem duplicate genes were identified relatively homogenously throughout the genomes of B73 and PH207 (fig. 2, supplementary fig. 3). A lower density of tandem duplicates was observed around the centromere where the density of genes is generally lower (Schnable et al. 2009). To test what variables most explained the distribution of tandem duplicates in the genome, a general linear model was fit with number of tandem duplicates per 1 Mb window regressed against number of genes, number of RNA transposable elements, number of DNA transposable elements, and subgenome within each window. With regards to subgenome, maize is a paleopolyploid that has returned to a diploid state. Two subgenomes remain in the diploid from the most recent allopolyploid event and have been previously characterized based on the number of retained co-orthologous genes to other grass species including sorghum and rice (Schnable et al. 2011; Brohammer et al. 2018). A number of differences are present between the subgenomes such as expression level (Schnable et al. 2011).

**Fig. 2.**
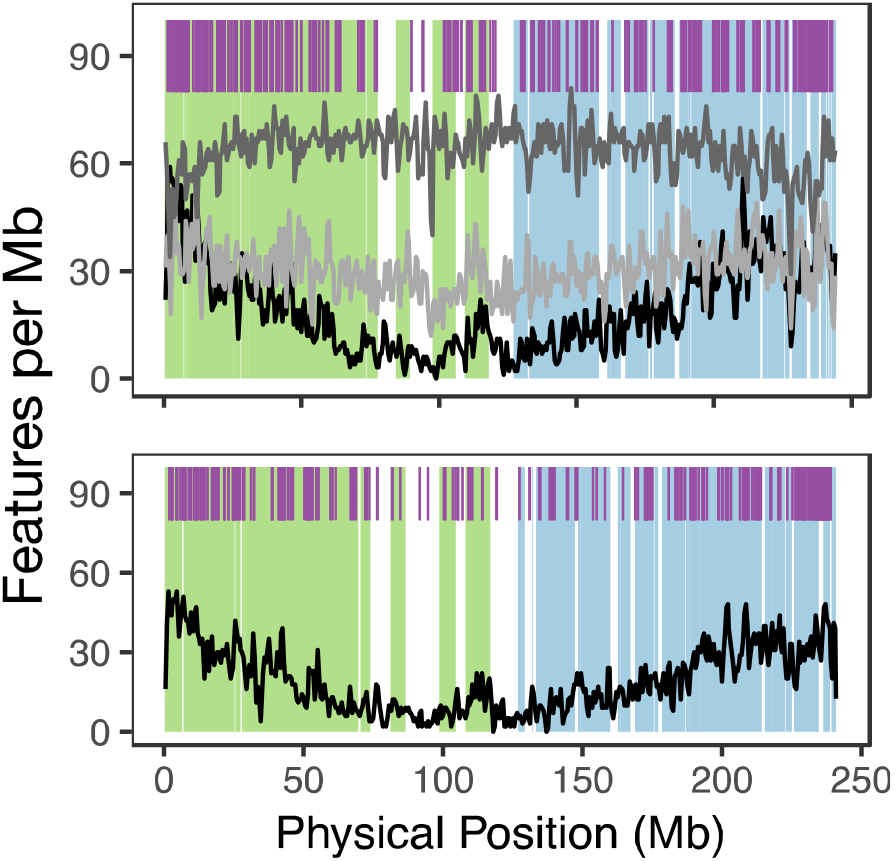
Genomic locations of maize tandem duplicates. Purple ticks show tandem duplications. Black line shows gene density, dark grey line shows RNA transposable element density, light grey line shows DNA transposable elements per Mb. Subgenome 1 is shown in green shading and subgenome 2 is shown in blue shading. The top panel shows B73 chromosome 2, and the bottom panel shows PH207 chromosome 2. All chromosomes can be found in supplementary fig. 3.

As expected, gene density explained the most variance in tandem duplicate density per window (t-test of regression coefficient, *p* < 0.001), and only 1% more variance was explained by a model containing all of these factors than a model with only gene density. When testing the effects of transposable elements, both class 1 and class 2 transposable element density were significant at the *p* < 0.05 threshold, and higher transposable element density was associated with higher tandem duplicate density. Within the general linear model, maize subgenome 2 was a significant negative factor while subgenome 1 was a significant positive factor (*p* < 0.001). Within tandem duplicate gene clusters, on average between the two genotypes 49.5% were in subgenome 1 (37.7% genome-wide), 21.6% were in subgenome 2 (24.0% genome-wide), and 28.9% were in non-syntenic regions (38.3% genome-wide) of the genome, consistent with the results of the general linear model where subgenome 1 has proportionally more tandem duplicates relative to the number of genes in the subgenome (table 1).

### Extant tandem duplicates date to two distinct periods

Phylogenetic analyses were used to estimate the date of origin of tandem duplicates. For each tandem duplicate cluster, all maize B73 and PH207 homeologs (i.e. subgenome 1 and subgenome 2 copies) and the corresponding sorghum gene for each of the tandem duplicate clusters were included. These phylogenetic trees were calibrated based on the estimated divergence time of maize and sorghum at approximately 12 million years ago (Swigonová et al. 2004). Our hypothesis was that tandem duplicate gene clusters that were shared between B73 and PH207 would be older than those that are unique to either one of the genomes. Indeed, we see examples of tandem duplicates that were shared and date near the divergence time of maize and sorghum at approximately 11.7 million years ago (fig. 3A). Conversely, fig. 3B shows an example of a private tandem duplicate that is only in B73 and arose approximately 33,000 years ago.

**Fig. 3.**
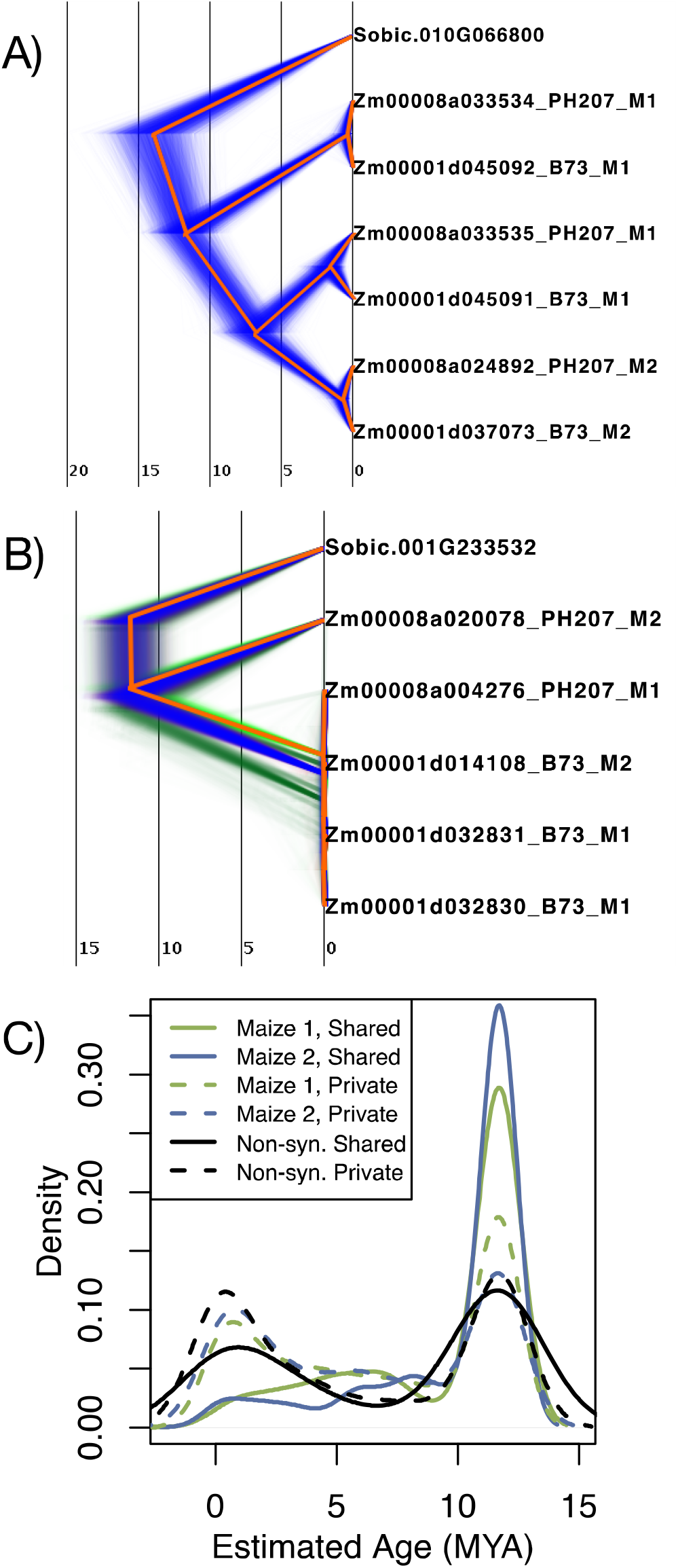
Date estimates of maize tandem duplications. A) Example of a BEAST tree that dated a tandem duplicate gene cluster as an ancient duplication. B) Example of a BEAST tree that dated a tandem duplicate gene cluster as a relatively recent duplication. In both trees red shows the consensus topology and alternate shading color indicates alternate topologies. C) Distribution of estimated syntenic and non-syntenic duplication ages. Shared tandem duplicate clusters are contained in both B73 and PH207 and private are only duplicated in one of the two genomes.

Across all the tandem duplicate gene clusters a bimodal age distribution was observed, with most tandem duplicates either dating to approximately the time of maize and sorghum divergence or dating quite recently in evolutionary time (fig. 3C). Consistent with our hypothesis, tandem duplicate gene clusters that were shared between the two genomes were almost exclusively in the peak of ancient tandem duplicates for both subgenome 1 and subgenome 2 gene clusters. In addition to being present in the ancient peak, a large number of the private syntenic duplicates also arose relatively recently. Of the 1,044 private clusters, 628 had an estimated date in the ancient peak and likely represent a gene loss event in one genotype and not the other. A comparable number of non-syntenic tandem duplicate gene clusters arose during both of the bimodal age peaks, similar to what was observed for private syntenic tandem duplicate gene clusters.

One explanation for the large proportion of inferred recent duplications is the action of gene conversion. Gene conversion would cause tandem duplicate genes to have higher sequence similarity than non-recombining duplicates of the same age, and thus would bias estimates toward recent events. Gene conversion is often associated with increased GC content due to GC-biased gene conversion (Pessia et al. 2012). Indeed, tandem duplicate genes showed a higher GC content on average than the GC content that was observed for all maize genes whether in the syntenic or non-syntenic portion of the genome (fig. 4A). If GC biased gene conversion were contributing to the peak of tandem duplicates that date relatively recently (less than 2 million years ago), we would expected that tandem duplicates within the young peak would have higher GC content than tandem duplicates within the old peak. However, the opposite was actually observed. Tandem duplicates that dated in the older peak had consistently higher GC content, and those that dated in the younger peak exhibited a high proportion of low GC content genes (fig. 4B). Thus, GC-biased gene conversion was likely not artificially deflating age estimates between duplicates to a substantial degree.

**Fig. 4.**
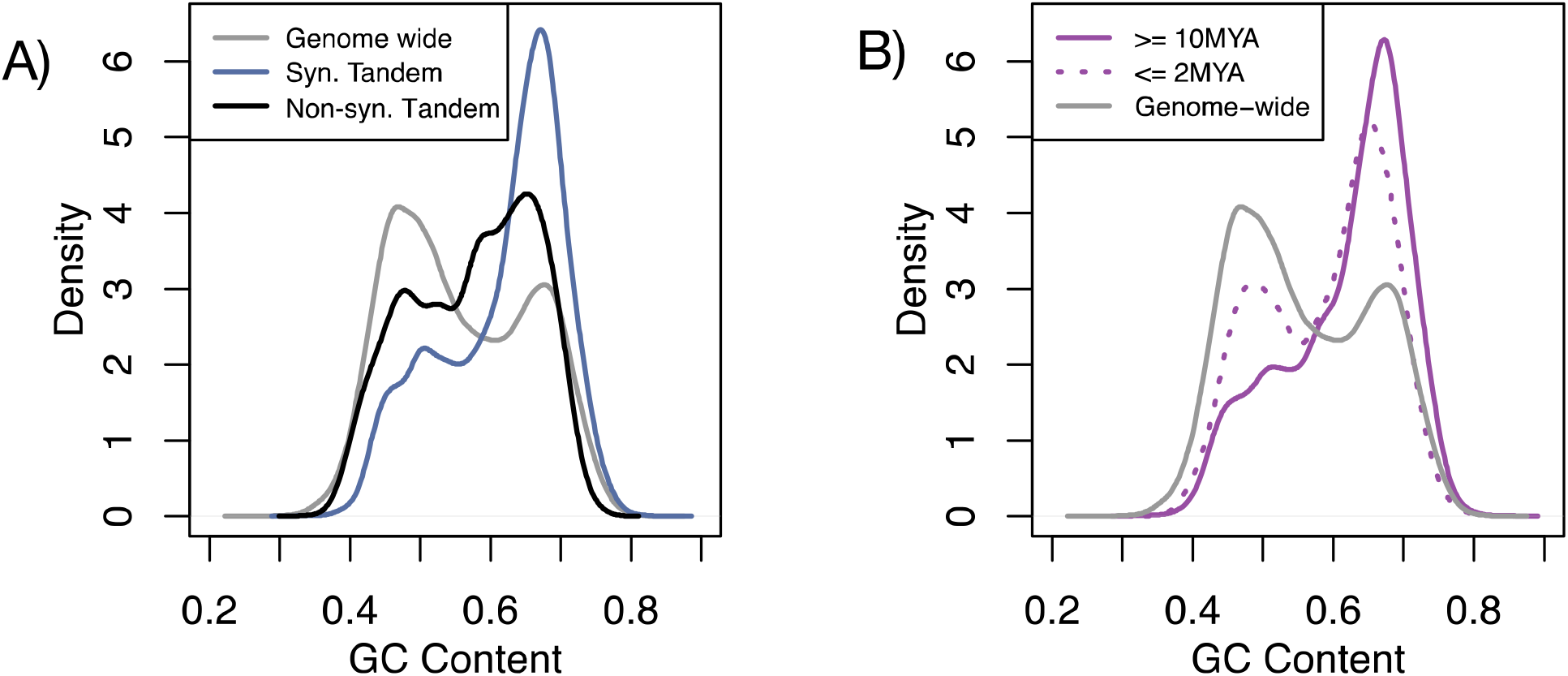
GC content of tandem duplicate gene clusters. A) GC content of genome wide genes (grey), syntenic tandem duplicates (blue), and non-syntenic tandem duplicates (black). B) GC content of ancient (>= 10MYA, purple solid) and recent (<= 2MYA, purple dotted) tandem duplications. Distributions contain both B73 and PH207 tandem duplicate gene clusters.

### Tandem duplicated genes are shorter and more likely to contain LTR transposable elements

One possible mechanism through which tandem duplicates could arise is through transposable elements. Some TIR elements are enriched for local movement and could contribute to tandem duplication of genes that are captured and moved locally. In contrast, LTR elements do not typically move locally, but if they were to randomly insert in a position of local proximity we would expect to observe a higher proportion of single exon genes in tandem duplicate gene clusters. In both B73 and PH207 a higher proportion of single exon genes in tandem duplicate gene clusters than was observed genome-wide (fig. 5 A and B). Instances in which a tandem duplicate cluster comprised a gene with multiple exons and its tandem duplicate contained only a single exon would further support the mechanism that the gene was duplicated through an RNA intermediate. Of the clusters that had a single exon gene (24% of total clusters), only one-third in B73 and two-thirds in PH207 also had a gene with multiple exons. However, 835 (B73) and 815 (PH207) clusters had multiple exons in both genes in the tandem duplicate gene cluster. These results indicate that while some tandem duplicate genes may have arisen through an RNA intermediate, this was not the only or even predominant mechanism.

**Fig. 5.**
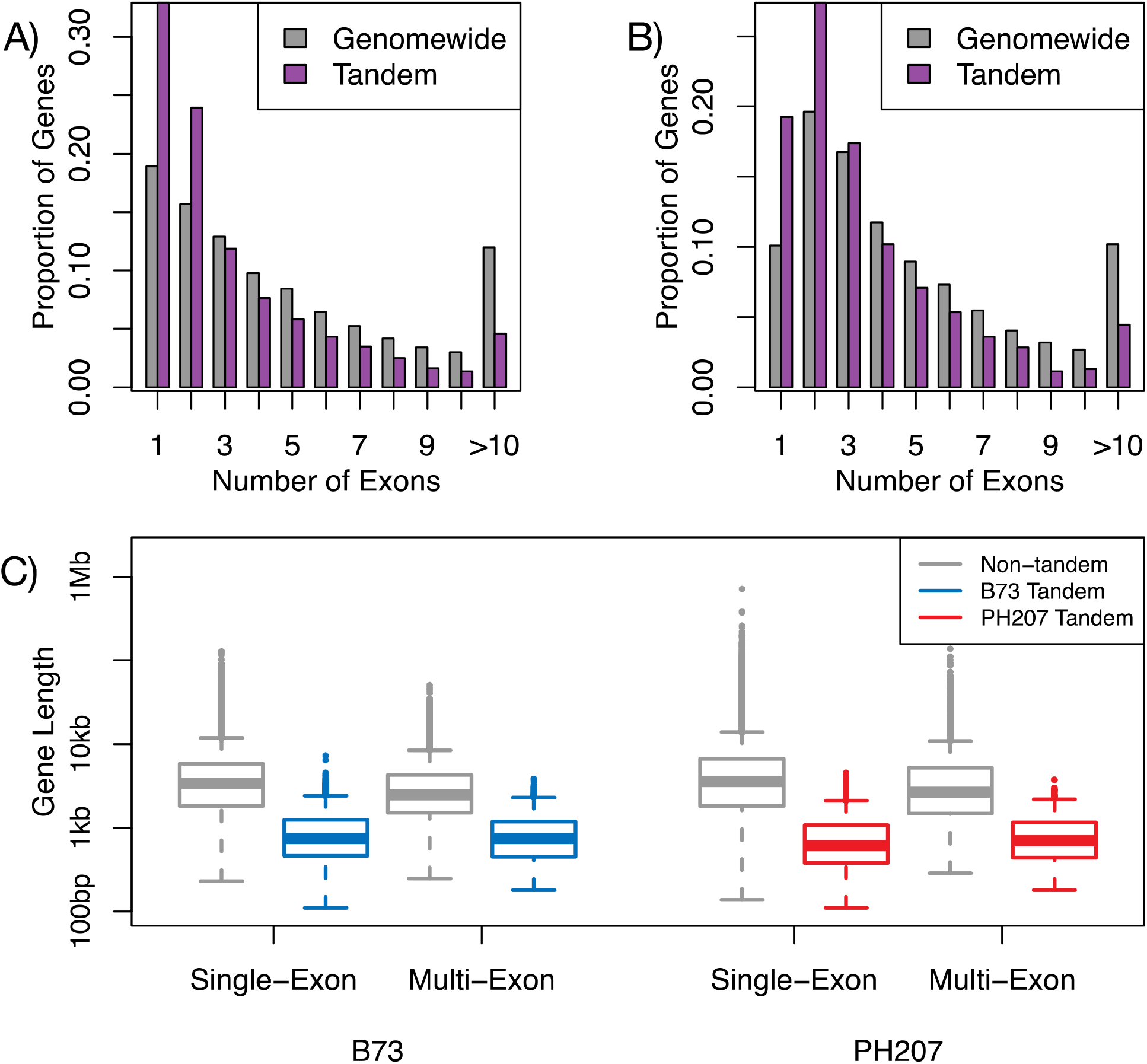
Size distribution of tandem duplicate gene clusters. A) Number of exons in B73 tandem duplicates relative to all other genes genome-wide. B) Number of exons in PH207 tandem duplicates relative to all other genes genome-wide. C) Gene length distribution for single-exon genes and multi-exon genes in B73 and PH207.

Another explanation for having a higher than expected rate of single exon genes in tandem duplicate gene clusters is if these genes were generally shorter and therefore easier to copy. For both single exon genes as well as genes with multiple exons in both B73 and PH207, genes that were in tandem duplicate gene clusters were shorter than the genome-wide distribution of gene sizes for single and multi exon gene models based on full gene model length (fig. 5C).

In addition to physically copying genes, transposable elements can also contribute to generating tandem duplicate gene clusters by providing microhomology for nonhomologous recombination. We investigated the relative proximity of tandem duplicate gene clusters to LTR, LINE, SINE, and TIR elements and found no difference in distance to nearest transposable element for tandem duplicate genes versus the genome-wide distribution (supplementary fig. 4). However, there was a substantial enrichment of LTRs inserted into tandem duplicate genes (20.0% of tandem duplicates contained LTRs versus 8.2% of non-tandem duplicates) and a de-enrichment of tandem duplicate genes that were captured (the entire genic sequence being nested within an element) by LTRs relative to the rate in non-tandem duplicates (2.6% of tandem duplicates versus 4.6% of non-tandem duplicates; table 2).

**Table 2.** Counts of tandem duplicate genes and non tandem duplicate genes that contain a transposable element insertion or are nested within a transposable element in the B73 genome.

| TE Class | Tandem Gene Contains | Tandem Gene Captured | Non-Tandem Contains | Non-Tandem Captured |
| --- | --- | --- | --- | --- |
| LTR | 889 | 115 | 2861 | 1598 |
| LINE | 0 | 1 | 4 | 16 |
| SINE | 0 | 10 | 0 | 166 |
| TIR | 0 | 127 | 48 | 1807 |

### Recent tandem duplicates evolve at different rates than other maize genes

We were interested in examining the relative substitution rates of recent tandem duplicates to infer potential evolutionary trajectories of novel tandem duplications. Substitution rates of recent tandem duplicate genes present in syntenic regions were compared to non-tandem duplicate maize genes and grass orthologs with Clade models in PAML (Yang 2007; Weadick and Chang 2012). Only duplications that were private to a subgenome and private to either B73 or PH207 were analyzed. A total of 120 grass orthologous groups with maize tandem duplicates met these filtering criteria (see Methods). Four competing hypothesis were tested that included testing 1) grass genes equal maize genes equal tandem duplicate genes evolutionary rates, 2) grass genes do not equal maize genes but equal tandem duplicate genes evolutionary rates, 3) grass genes equal maize genes do not equal tandem duplicate genes evolutionary rates, and 4) grass genes do not equal maize genes do not equal tandem duplicate genes evolutionary rates. The majority (74.2%) of the tandem duplications did not show evidence of evolving at a different rate from their grass orthologues (test 1 and 2). Of the 31 (25.8%) tandem duplicate clusters that have evolved at a different rate than the remainder of the tree (test 3 and 4), 10 tandem duplicates showed lower dN/dS than the remainder of the tree, seven showed higher dN/dS than non tandem duplicates, and 14 did not have enough dS to compare substitution rates (supplementary fig. 5). It should be noted that these are all coding sequence based tests and do not assay the promoter sequences which could alter gene functionality. Additionally, specific gene conversion events could potentially alter dN and/or dS which could impact omega estimates.

## DISCUSSION

Tandem duplicate gene studies in plants have primarily focused on single loci and questions about evolutionary mechanism and functional impacts on a genome-wide scale have been limited by available genomic resources. Maize offers a unique opportunity to address questions regarding tandem duplicate origin and evolution given its large, large highly repetitive genome and the availability of multiple high quality and well-annotated *de novo* genome assemblies. Using these resources, we showed that tandem duplicate gene clusters are prevalent in maize, and there is variation within maize lines for tandem duplicate content in the genome. While tandem duplicate clusters are dispersed in genome space, standing variation in tandem duplicates date to two distinct times. A variety of features in the genome were evaluated for association with tandem duplicates. Tandem duplicate genes are shorter than non-tandem duplicate genes and are more likely to contain LTR transposable elements, which may speak to their origin. For a subset of the tandem duplicates that could be tested, approximately one-quarter were evolving at different rates than other maize genes. These duplicates along with others that are likely evolving in regulatory control have potential to generate new functional variation.

While tandem duplicate gene clusters are abundant and generally dispersed throughout the genome at a density similar to the genome-wide gene density, we did observe some bias in the location of tandem duplicate gene clusters. Specifically, maize subgenome 2 has proportionally fewer tandem duplicates than either subgenome 1 or non-syntenic regions relative the gene densities. There are several explanations for this. One is that duplications do not arise in maize subgenome 2 as readily as other genomic regions. This is unlikely, however, because the patterns of transposable elements and recombination dynamics of subgenome 2 are similar to subgenome 1 (Schnable et al. 2011). Additionally, subgenome 2 is under weaker purifying selection than subgenome 1 (Schnable et al. 2011), meaning that gene duplications should be more prevalent in subgenome 2 but this was not observed. Another explanation is that genes in subgenome 2 “degenerate” more rapidly than in subgenome 1, and that genes that are truly tandem duplicates are too divergent at the sequence level at this point in time to be identified as duplicates by our methods. It has also been hypothesized that subgenome 2 has a higher ongoing deletion rate than subgenome 1 and may have proportionally fewer tandems due to a faster rate of deletion from the genome in subgenome 2.

While dispersed throughout the genome, the estimated ages of standing tandem duplicates is not dispersed throughout evolutionary time. In fact, tandem duplicates have a bimodal distribution of estimated ages (fig. 3C). This suggests that tandem gene duplication has occurred in bursts throughout maize evolutionary history. The older peak in the estimated age distribution coincides with the divergence of maize from sorghum (Swigonová et al. 2004). This is not unexpected, because the genomic instability and rearrangements caused by an allopolyploidy event can result in many tandem duplicates (Blanc and Wolfe 2004). The more recent peak coincides with the expansion of maize as an agricultural plant (Wang et al. 2017). Strong selection for agronomic traits and rapid increase in population size may reduce the efficacy of purifying selection against recent tandem duplications.

In addition to determining when in evolutionary history tandem duplicates arose, we also tried to determine their mechanistic origin. Our results suggest that tandem duplicates have a close association with transposable elements. This is evident at both the genomic distribution level (fig. 2), and direct comparison of gene model annotations and transposable element annotations, where there is an enrichment of LTRs that are nested within tandem duplicates relative to non-tandem duplicates (table 2). This suggests that tandem duplicates may arise through a mechanism that preferentially operates on highly repetitive sequence such as transposable elements. One such mechanism is unequal crossing over (Smith 1976). Errors in meiotic chromosome pairing are often the result of repetitive elements like tandemly arrayed genes (Yandeau-Nelson et al. 2006), and may contribute to the significant level of gene copy number variants observed in maize (Springer et al. 2009; Swanson-Wagner et al. 2010). Tandem duplicates may also be generated via transcription-mediated mechanisms associated with RNA transposable elements. We observed a higher proportion of single exon genes in tandem duplicate gene clusters relative to non-duplicated genes. A strong signature of an RNA-intermediate in a tandem duplication would be if one gene within the tandem duplicate contained multiple exons and the other gene within the tandem duplicate contained only a single exon that is the product of a spliced mature mRNA. We did not see this pattern in many of tandem duplicate gene clusters. Tandem duplicate genes, whether single exon or multi exon, however, were generally shorter than non-tandem duplicates. This points to an explanation that single exon genes, which are generally shorter, are easier to copy intact through mechanisms such as non-homologous recombination (Smith 1976), rather than a RNA intermediate.

In this study, we constrained our analyses to annotated genes. However, tandem duplication can affect regulatory elements or gene fragments (c.f. Rogers et al. 2017). Duplication of functional elements that are not full-length protein coding genes likely has an impact on phenotypic variation, and therefore, evolution of genome structure. Our work presents a special case of tandem duplications, in which entire genes are duplicated. However, we show that even this special case of tandem duplication can affect thousands of genes genome-wide and has the potential for functional outcomes.

## MATERIALS AND METHODS

### Tandem Duplicate Identification

Putative tandem duplicate clusters were identified by comparing the B73 version 4 (Jiao et al. 2017) and PH207 version 1 (Hirsch et al. 2016) maize genome assemblies each to the Sorghum bicolor v3.1 genome assembly (DOE-JGI, http://phytozome.jgi.doe.gov/) with SynMap v2 (Lyons et al. 2008) as previously described (Brohammer et al. 2018). Using these methods genes could be defined as tandem duplicates with up to 15 intervening genes. To remove any false positive assignments from SynMap and identify any false negative assignments, the longest transcripts from adjacent maize genes were translated to amino acids, aligned (10 interactions of refinement) with Clustal-omega (Sievers et al. 2011), and back-translated to nucleotides. Pairwise similarity, was calculated with the “compute” program from the “analysis” package (Thornton 2003). The alignment similarity was down-weighted for the proportion of gaps open during alignment by calculating the similarity in aligned regions multiplied by the proportion of the total alignment not in gaps. A distribution of adjusted pairwise similarities for adjacent genes in the B73 and PH207 assemblies is shown in supplementary fig. 6. Adjacent genes from this method and any additional tandem duplicate genes clusters with intervening genes identified using SynMap with at least a 0.3 adjusted pairwise similarity were retained for downstream analyses.

### Tandem Duplicate Gene Cassette Identification

Tandem duplicate cassettes were identified by comparing annotated gene coordinates among individual tandem duplicate clusters. A schematic of the procedure for identifying cassette duplications is shown in supplementary fig. 2. Genes within each tandem duplicate cluster were sorted from lowest coordinate to highest coordinate. Tandem duplicate clusters on the same chromosome were sorted by the start coordinate of the first gene. Tandem duplicate clusters that overlapped each other were identified. Clusters that were fully nested within another cluster were not considered as cassettes. Tandem duplicate clusters that were interleaved within each other were retained as putative cassette duplications (supplementary fig. 2). **General Linear Model Analysis of Factors Explaining Tandem Duplicate Gene Density** A general linear model was fit to explain variation in tandem duplicate gene density within the R version 3.4.1 computing environment (Team 2017). The model regressed tandem duplicate gene density against all annotated gene density, RNA transposable element density, DNA transposable element density, and subgenome assignment. All density calculations were performed in 1Mb windows across the genome. We defined density as the proportion of bases in each window that were contained in tandem duplicate genes, all annotated genes, RNA transposable elements, or DNA transposable element. Gene and transposable element annotations were obtained from Gramene (ftp://ftp.gramene.org/pub/gramene/release-55/gff3/zea_mays/). Subgenome assignments were from previously reported syntenic block assignments (Brohammer et al. 2018). Because annotated gene density is correlated with subgenome assignment (Schnable et al. 2012), we tested models with and without an interaction between subgenome and gene density. Models with an interaction between subgenome and gene density did not significantly improve model fit (ANOVA of nested models, p> 0.2).

### Duplication Date Estimation

The dates of tandem duplications were estimated with BEAST version 2.4.7 (Bouckaert et al. 2014). Amino acid alignments of the subgenome homeologues, tandem duplicates, and putative Sorghum ancestral genes that were previously determined (Brohammer et al. 2018) were generated with Clustal-omega version 1.2.1 (Sievers et al. 2011). The alignments were back-translated to nucleotides. Each gene alignment was analyzed with BEAST with the following parameters: a GTR+Gamma nucleotide substitution model, estimated transition probabilities and equilibrium base frequencies, a random local clock to allow for branch-specific rate variation, and a monophyletic divergence between the maize subgenomes with a prior of ~N(11.9,1) on the divergence date. The MCMC was run for 10,000,000 steps.

Resulting trees from the BEAST analysis were parsed to obtain the time to most recent common ancestor (TMRCA) between tandem duplicate genes in both B73 and PH207 to avoid double counting shared duplications. Gene duplications likely do not follow the infinite sites mutational model (Kimura 1969; Ohno 1970), and identity by state does not necessarily imply identity by descent. This analysis assumed that tandem duplicates are evolving along truly separate trajectories, and that gene conversion among tandem duplicates is negligible.

### Intersection of Tandem Duplicates and Transposable Elements

Structural annotation of transposable elements in B73 was obtained from Gramene (ftp://ftp.gramene.org/pub/gramene/release-55/gff3/zea_mays/repeat_annotation/). Genes containing transposable element insertions and genes captured by transposable elements were identified using ‘bedtools intersect’ (Quinlan and Hall 2010) requiring an overlap fraction of 1.0.

### Relative Rates Calculations

Relative rates of sequence evolution of maize tandem duplicates were compared with other grass genes by performing Clade model C (CMC) tests (Weadick and Chang 2012) in PAML v4.9e (Yang 2007) on tandem duplicates in orthologous gene groups. Orthologous gene groups were identified among publicly available grass genomes from Phytozome V12 and Ensembl Plants V34 using OrthoFinder v1.14 (Emms and Kelly 2015). The species, sources, and versions of the genomes used as OrthoFinder input are shown in supplementary table 2. OrthoFinder was run with the “dendroblast” orthologue search method, and default parameters for MCL clustering.

Amino acid sequences from B73 and PH207 were kept in separate files to allow them to be compared to each other as well as to other grasses. Only orthologous gene groups between 10 and 75 genes were retained for analysis because larger orthologous groups are likely to be gene families that have already diverged in function, and smaller groups do not have enough genes to test relative rates. A distribution of orthologous gene group sizes is shown in supplementary fig. 7. Orthologous gene groups that contained between 10 and 75 genes, contained maize tandem duplicates, and contained complete tandem duplicate clusters (i.e., tandem duplicate clusters that were not split among multiple orthologous groups) were retained for downstream analysis.

Within each of the orthologous groups that passed the above filtering, the amino acid sequences were aligned with clustal-omega and then back-translate to nucleotides using the CDS sequences. Alignments were filtered to contain only sites with at most 50% gaps across all sequences, because gaps greatly increased computation time and were not informative for calculating substitution rates. Maximum likelihood trees were estimated from the filtered alignments with RAxML, using the default rapid hill-climbing search algorithm and a GTR+Gamma nucleotide substitution model.

Four models were fit to the filtered alignments and trees with the ‘codem? program in PAML (supplementary fig. 5). We tested whether certain foreground branches of the tree exhibited significantly divergent evolutionary rates relative to the remainder of the tree. Model 1 marks maize genes and common ancestors of maize genes as different from other grass genes. Model 2 marks maize tandem duplicates and common ancestors of maize tandem duplicates as different from all other genes. Model 3 distinguishes maize tandem duplicates and common ancestors of maize tandem duplicates from maize genes, and Model 4 sets maize genes as different from other grass genes (supplementary fig. 5). The null model treated every clade as evolving under the same constraint (Weadick and Chang 2012). The best-fitting model for each orthologous group was identified via a likelihood ratio test against the null model. In orthologous groups where maize genes were evolving at a different rate than tandem duplicate genes, omega was compared between tandem and non-tandem maize genes to determine relative constraint. If omega in tandem duplicates was higher than non-tandem duplicates, the tandem duplicate was classified as under weaker constraint, and if it was lower than the nontandem duplicate it was classified as under stronger constraint. If omega was larger than 10 no test was done as dS was considered too small to test.

### Code Availability

Scripts to perform tandem duplicate identification, sequence alignment and back-translation, orthologous gene group identification, and relative rates comparisons are available at https://github.com/TomJKono/Maize_Tandem_Evolution.

## ACKNOWLEDGMENTS

This work was supported by the National Science Foundation Plant Genome Research Project grant (IOS-1546727) to CNH and SEM. ABB was supported by a DuPont Pioneer Bill Kuhn Honorary Fellowship and a MnDRIVE Global Food Ventures fellowship. The authors acknowledge the Minnesota Supercomputing Institute (MSI) at the University of Minnesota for providing resources that contributed to the research results reported within this paper http://www.msi.umn.edu.

